# IAMSAM : Image-based Analysis of Molecular signatures using the Segment-Anything Model

**DOI:** 10.1101/2023.05.25.542052

**Authors:** Dongjoo Lee, Jeongbin Park, Seungho Cook, Seongjin Yoo, Daeseung Lee, Hongyoon Choi

## Abstract

Spatial transcriptomics is a cutting-edge technique that combines gene expression data with spatial information, allowing researchers to study gene expression patterns within tissue architecture. Here, we present IAMSAM, a user-friendly web-based tool for analyzing spatial transcriptomics data focusing on morphological features. IAMSAM accurately segments tissue images using the Segment-anything model, allowing for the semi-automatic selection of regions of interest based on morphological signatures. Furthermore, IAMSAM provides downstream analysis, such as identifying differentially expressed genes, enrichment analysis, and cell type prediction within the selected regions. With its simple interface, IAMSAM empowers researchers to explore and interpret heterogeneous tissues in a streamlined manner.

## Background

Spatial transcriptomics (ST) enables the analysis of gene expression patterns inside tissues while maintaining their spatial context (1). However, researchers often encounter difficulties when working with ST data due to its complexity, high-dimensionality, spatial constraints, large data volumes, and the lack of user-friendly tools (1,2) For instance, clustering spots or cells within one or multiple ST libraries must exhibit spatial continuity for each cluster, which requires the use of complex algorithms (3). Furthermore, the manual process of identifying genes associated with specific regions, based on domain knowledge such as pathologist-labeled annotation, introduces a subjective analytic workflow (4,5). Integrating the interpretation of tissue image patterns, along with multidimensional molecular information, allows researchers to gain a deeper understanding of the pathophysiology within spatial contexts (6–8). Therefore, an interactive and user-friendly interface for ST to analyze tissue images should also be developed to facilitate improved communication for basic researchers, clinicians, and bioinformaticians.

Here, we introduce IAMSAM (Image-based Analysis of Molecular signatures using the Segment-Anything Model), a user-friendly web-based tool designed to comprehensively analyze ST data, enabling a better understanding of complex tissues by integrating images with molecular information. IAMSAM leverages the power of the Segment-anything model (SAM), a state-of-the-art deep learning model developed by Meta (9), to identify regions of interest (ROIs) from tissue images in ST datasets. The SAM model exhibits exceptional performance, achieving real-time performance and efficiently utilizing computational resources. Moreover, it stands as the first foundation model for general image segmentation, providing interactive prompting capabilities. It has been specifically designed to address the problem of zero-shot image segmentation pre-trained on an extensive and diverse dataset consisting of over 1 billion masks derived from 11 million images, ensuring its robust performance. We used this model to handle various tissue images (e.g., H&E, DAPI, and immunofluorescence images), taking advantage of its effectiveness and adaptability to handle different image distributions and workloads through zero-shot or few-shot learning. This excellency leads to conducting various downstream analyses such as identifying differentially expressed genes (DEGs), enrichment analysis, and cell type prediction of user-selected regions. In this study, we demonstrated the usage of IAMSAM with publicly available ST datasets. With its simple and accessible interface, IAMSAM enables researchers to explore and interpret their ST data user-friendly, which can lead to new insights into gene expression patterns associated with pathophysiology and potential biomarkers for diseases.

## Results

### Overview of IAMSAM

IAMSAM is a web-based tool designed for analyzing ST data, based on a general purpose image segmentation algorithm named ‘Segment-anything’ **(Figure 1)**. It utilizes the SAM for H&E image segmentation, which allows for morphological guidance in selecting ROIs for users. IAMSAM offers users with two modes for running the SAM algorithm: everything-mode and prompt-mode. In the everything-mode, users have the option to select one or multiple segments from the automatically generated segments. On the other hand, the prompt-mode allows users to draw rectangle boxes, which serve as input prompts for the SAM model. IAMSAM automatically extracts the gene expression profile from the chosen ROIs, identifying not only DEGs between the ROIs and non-ROIs but also enriched functional terms associated with these DEGs. Furthermore, IAMSAM provides cell type estimation of the selected regions, which can help users gain valuable insights into the cellular composition and heterogeneity of the tissue.

**Figure 1.**
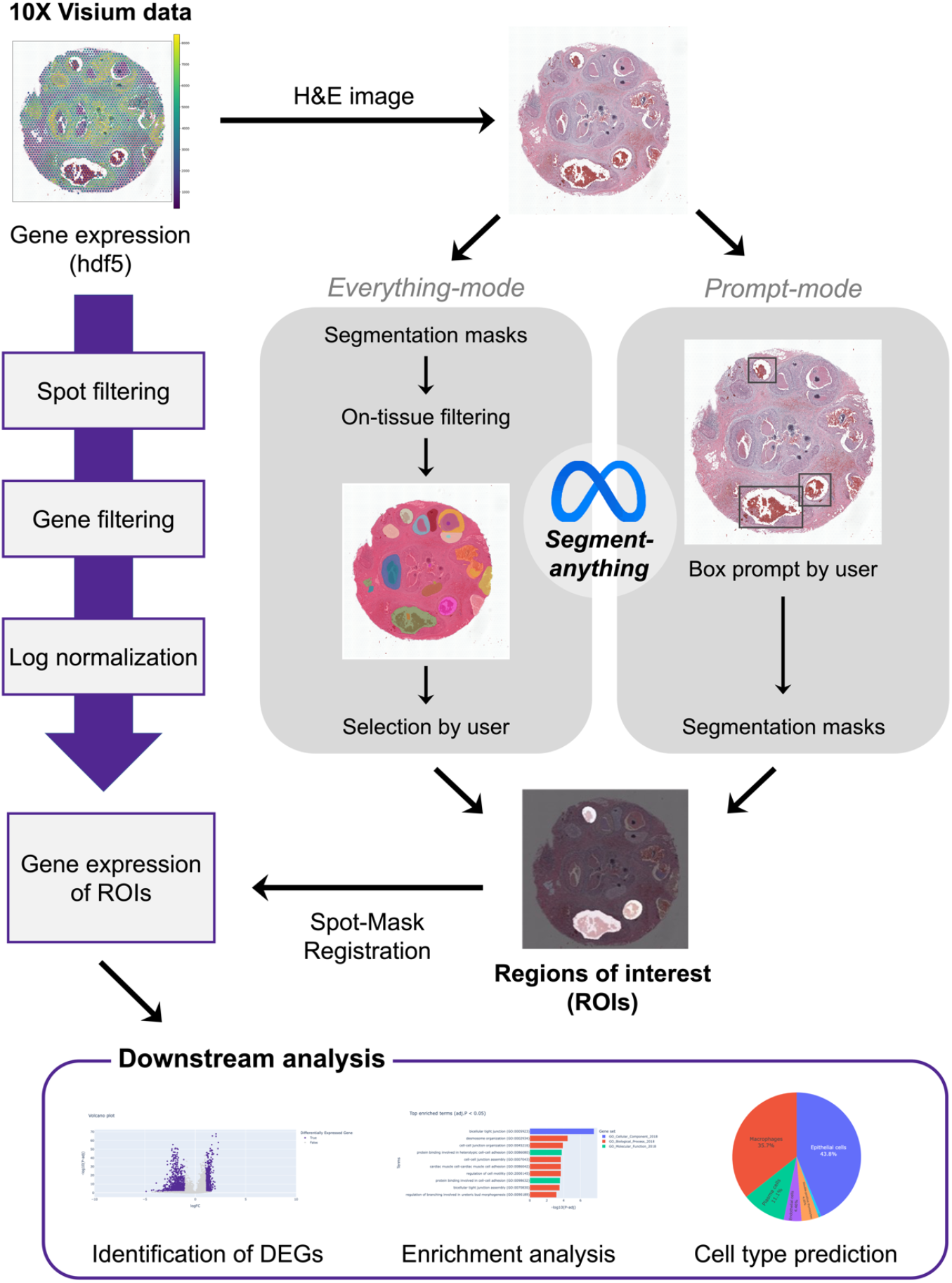
Workflow of IAMSAM. This figure provides an overview of the workflow of IAMSAM. The gene expression of ST data is preprocessed through spot filtering, gene filtering, and log-normalization step. The H&E image of the ST data is segmented using the segment-anything model in two different modes: everything-mode and prompt-mode. The selected ROIs are then subjected to downstream analysis, which includes DEG identification, enrichment analysis, and cell type prediction.

### H&E image segmentation

Hematoxylin and eosin (H&E) are widely employed to observe tissue structure, distinguish different histological features, and are considered a gold standard in the field of histopathology (10). Most ST platforms, particularly the 10x Visium platform, involve the inclusion of H&E staining and tissue imaging steps in the tissue preparation protocol (11). This unique feature of the Visium platform allows IAMSAM to utilize the H&E image. When users select the samples to analyze on the dropdown menu, the H&E image of the sample appears in the main visualization panel **(Figure 2a)**. After configuring multiple parameters, users can click the ‘Run SAM’ button to make inferences from the Segment-anything model (SAM). SAM takes the H&E slide images as input and creates a binary mask for each morphologically segmented region. IAMSAM visualizes these segment masks on the main visualization panel with a distinct palette, offering a user-friendly approach for researchers to analyze their ST data. Users can generate SAM masks and specify ROIs in two different modes, depending on their requirements or preferences. This approach not only reduces the time and effort required for manual annotation but also provides a more objective way of identifying morphological features and molecular signatures within the tissue.

**Figure 2.**
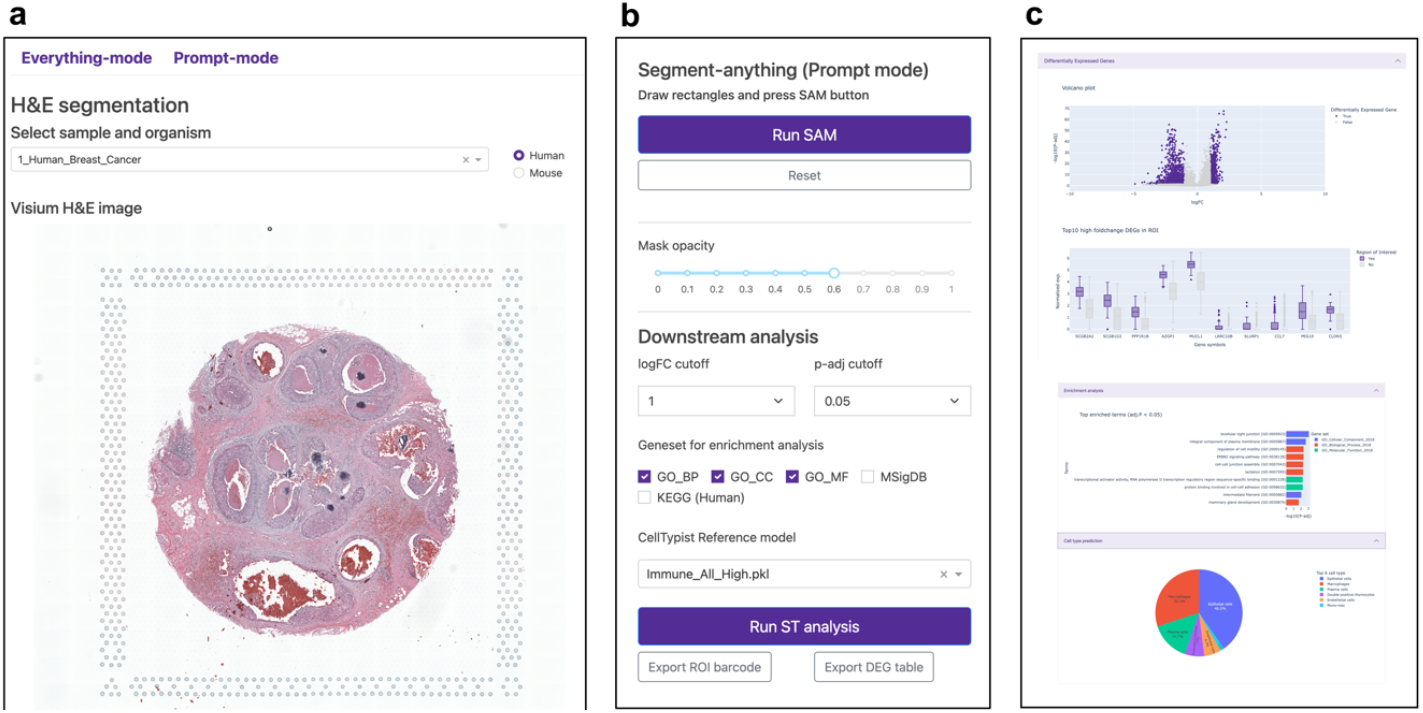
Three main panels in IAMSAM. This figure illustrates the three main panels in IAMSAM. (a) The main visualization panel displays the H&E slides of the ST data, along with the corresponding segmentation masks. These masks highlight different ROIs within the tissue image, allowing users visually explore and select specific ROIs. (b) The options panel provides a range of customization options for the analysis. It includes settings for the SAM, mask selection, and downstream analysis parameters. (c) The downstream analysis panel presents the analysis performed in IAMSAM, including DEG analysis, enrichment analysis, and cell type prediction.

### Everything-mode: automatic generation of masks in entire images

In the everything-mode, users can obtain segmented masks for the entire tissue image by simply clicking the “Run SAM” button. IAMSAM automatically segments the entire image, creating masks that distinguish various morphological features or regions within the tissue without requiring any additional prompts.

The ‘IOU score threshold’ parameter **(Figure 3a)** is a crucial factor for users to consider. The intersection-over-union (IOU) score is a metric used to measure the overlap between the predicted segmentation mask and the ground truth mask (9). By increasing the threshold value, the model becomes more stringent in accepting predicted masks. This means that only masks with a higher predicted IOU value, indicating better quality and accuracy, will be included in the final segmentation results. Consequently, the number of selected masks may decrease. Conversely, reducing the threshold makes the model more permissive in accepting masks, even if their predicted IOU is low. This relaxation of criteria can yield a higher number of masks, including those with potentially lower quality. Users should control the balance between the number of masks and their quality in the segmentation results, based on their specific requirements and preferences.

**Figure 3.**
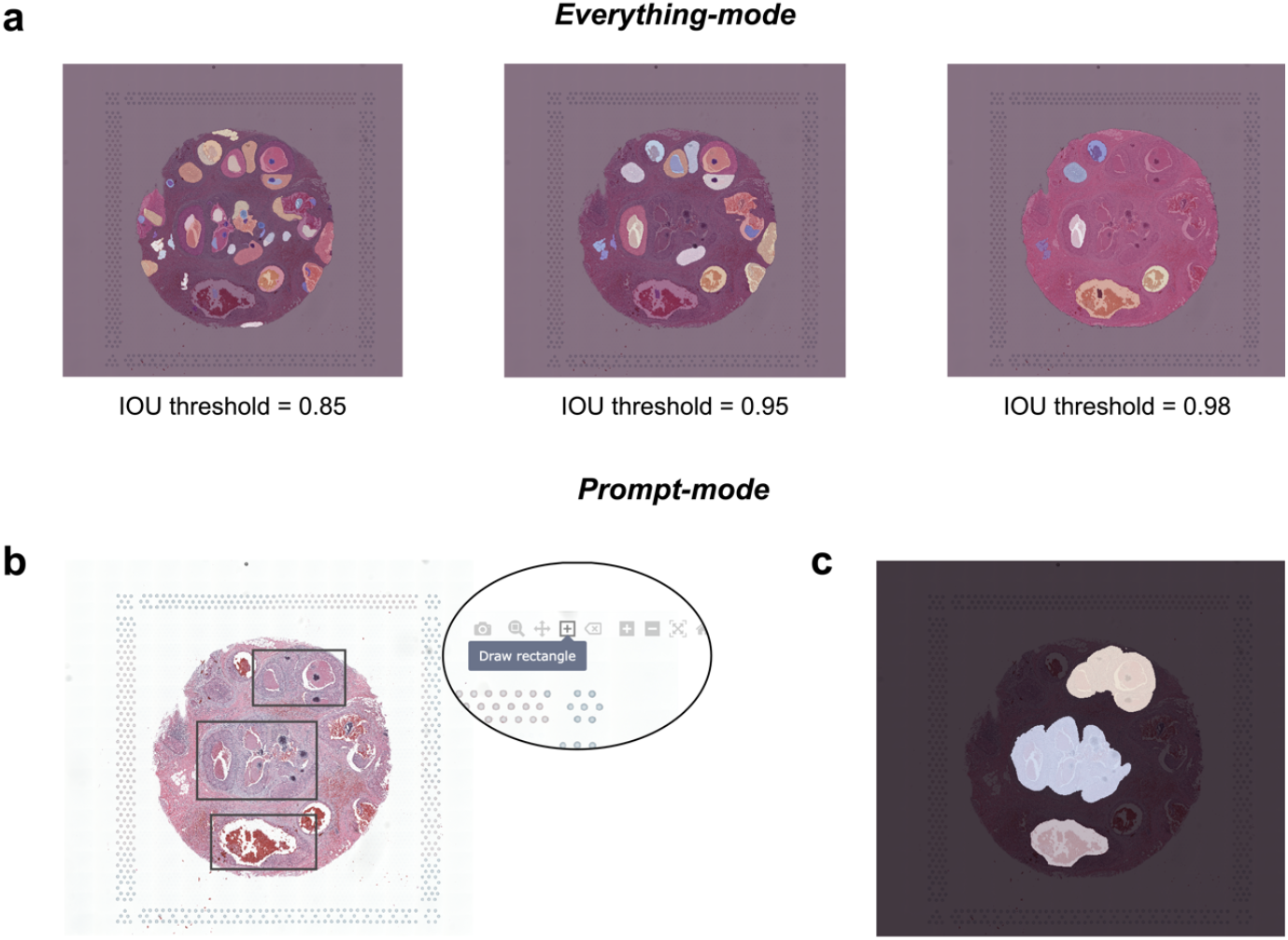
Everything-mode and prompt-mode. This figure introduces the two main modes of operation in IAMSAM: everything-mode and prompt-mode. (a) In everything-mode, IAMSAM generates segmentation masks for the entire tissue images. The IOU threshold directly affects the segmentation result, where a higher threshold leads to more precise segmentation but fewer selected masks. (b) In prompt-mode, users can provide prompts to the SAM model by drawing rectangle boxes on the visualization panel using the drawing tool provided by Plotly. When users input three rectangle boxes as drawn, IAMSAM returns the corresponding ROIs, as shown in (c).

After the segmentation, masks without containing any spots are filtered out, and the remaining masks are numbered in descending order based on their respective areas. Users can choose the mask number from a dropdown menu to select a particular mask. Alternatively, users can also just click on the masks directly in the main visualization panel. This feature is enabled through the interactive interface of Plotly (12), which allows the user to visualize the segmented regions and select the ROIs with ease. For an improved user experience, we also have added a feature that allows users to deselect a selected mask by simply re-clicking on it. After all, ROIs have been chosen, users can run downstream analysis on the ROIs with the ‘Run ST Analysis’ button. By enabling users to select the masks of interest through a simple click, IAMSAM streamlines the analysis of ST data and allows researchers to quickly identify relevant cell types and gene expression patterns in their samples.

### Prompt-mode: guided segmentation with box prompt

IAMSAM offers another mode called prompt-mode, which provides users with the flexibility to manually define the desired segments using rectangle boxes. This mode utilizes the prompt-input method of the original SAM algorithm, allowing users to specify boxes in the image that correspond to the objects they want to segment. Before running SAM, users can easily draw rectangles on the main visualization panel using the default rectangle drawing tool **(Figure 3b)**. Users can also conveniently track the number of boxes added and have the button to reset if any mistakes are made. Since box prompts are available in advance before running SAM, IAMSAM can run SAM in a batched manner, generating corresponding masks for multiple boxes simultaneously **(Figure 3c)**. If needed, users can utilize the zoom feature provided by Plotly when selecting ROIs in the prompt-mode. Upon clicking ‘Run SAM’, one or more masks are interpreted as the user’s areas of interest, and subsequent downstream analysis is performed in the same way as the everything-mode.

### Downstream analysis

The following downstream analysis consists of three panels: identifying DEGs, enrichment analysis (also known as geneset analysis), and cell type prediction **(Figure 2c)**. As all the downstream plots are interactively made, the various convenient features including auto-scaling, manual scaling, zoom-in, zoom-out, capture, and the management of the coordinates are supported for each plot.

The first panel is the DEG module, which includes both the volcano plot and the box plot. The volcano plot represents the log-fold change on the x-axis, where positive values indicate up-regulation in the ROIs, and the statistical significance on the y-axis. Users can set the ‘logFC cutoff’ and ‘p-adj cutoff’ in the parameter panel **(Figure 2b)**. Genes that meet the criteria of having a fold change value exceeding the FC cutoff and an adjusted p-value less than the adjusted p-value cutoff are displayed in purple, while the remaining genes are shown in gray. The box plot, on the other hand, focuses on the top 10 genes selected from the up-regulated DEGs within the ROIs. These genes are ranked based on their fold changes, reflecting the relative difference in expression levels between the ROIs and non-ROIs.

In the second panel, IAMSAM performs over-representation analysis (ORA) on the DEGs identified in the selected ROIs. The goal of ORA is to assess whether specific gene sets or functional categories are overrepresented among the DEGs, indicating their potential involvement in specific biological processes or molecular functions. IAMSAM offers users a choice of gene sets for enrichment analysis, including three GO (Gene Ontology) terms (biological process, cellular component, and molecular function), as well as gene sets from MSigDB (Molecular Signatures Database) and KEGG (Kyoto Encyclopedia of Genes and Genomes). Users can select the gene sets of interest based on their preferences to perform the enrichment analysis. IAMSAM calculates the statistical significance of the enrichment terms and filters them based on adjusted p-values. Only the terms that demonstrate statistical significance, with adjusted p-values below 0.05, are displayed in the form of a bar plot. This visualization allows users to easily identify the enriched terms and gain insights into the functional annotations associated with the DEGs.

For the last panel, IAMSAM provides cell type prediction within the selected ROIs. To perform cell type prediction, IAMSAM utilizes the CellTypist API (13), chosen for its high speed (14). Users can select from 19 reference models provided by CellTypist to ensure accurate predictions for their specific samples. Users need to choose the appropriate reference model that matches their sample. Additionally, IAMSAM automatically convert gene symbol between human and mouse, when the species of the sample and the reference model differ. The predicted proportions of cell types are visualized as a pie chart, displaying the top 6 cell types with the highest proportions for the sake of clarity and simplicity. This concise representation offers a clear overview of the predominant cell types present in the tissue sample and aids in understanding the cellular composition within the spatial context.

### Example usages of IAMSAM

We examined six different usage examples from publicly available datasets. These datasets were named as follows: Human_Breast_Cancer (15), Mouse_Colon (16), Mouse_Brain_H&E (17), Mouse_4T1 (18), Human_Prostate_Cancer (19), and Mouse_Brain_FL (20) **(Figure 4)**. Here, we utilized everything-mode for the former three samples and prompt-mode for the following two samples. The last sample was introduced for both applications to demonstrate the applicability of SAM to fluorescence images.

**Figure 4.**
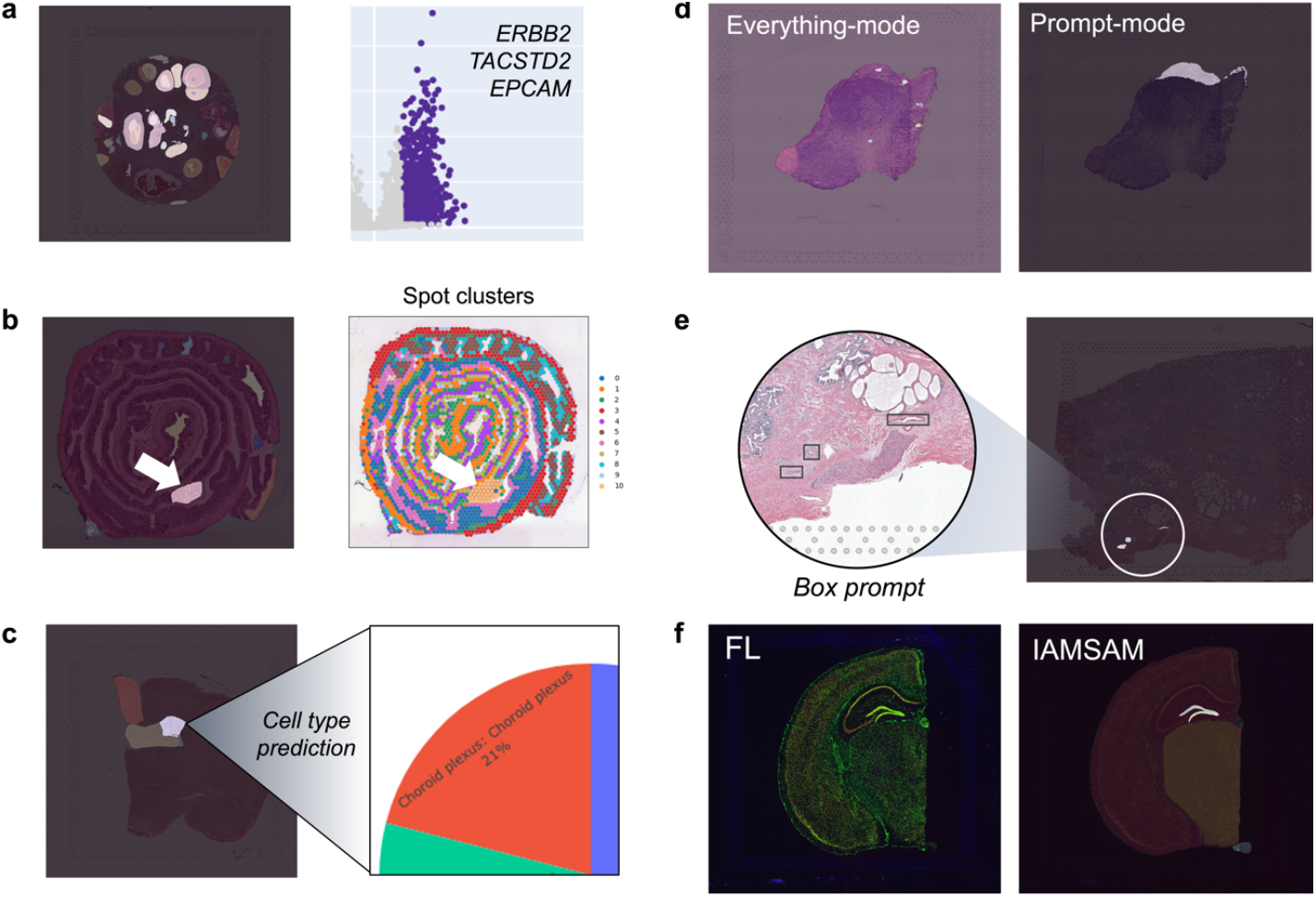
Example usage of IAMSAM. This figure presents six cases demonstrating the capability of IAMSAM. (a) IAMSAM’s everything-mode accurately identifies breast cancer marker genes as DEGs. (b) The segmentation results obtained from IAMSAM in the Mouse_Colon sample exhibit strong agreement with the results of spot clustering analysis, indicating consistent and reliable performance. (c) The application of IAMSAM’s everything-mode to the Mouse_Brain_H&E sample successfully reveals the presence of the choroid plexus (CP) structure, along with the co-occurrence of cycling microglia. (d) In the Mouse_4T1 sample, prompt-mode, rather than everything-mode, effectively identifies a stromal region located at the periphery of the tumor section. (e) By combining the zoom-in interface with prompt-mode, IAMSAM allows for the detailed examination of microscopic histology features, enhancing the analysis capabilities. (f) IAMSAM can be applied not only to H&E images but also to various imaging modalities, including fluorescence imaging, thereby expanding its utility in different experimental settings.

In the case of the Human_Breast_Cancer sample, our user-friendly interface enables easy selection of cancer-rich regions **(Figure 4a; Supplementary Figure 1)**. We observed that the selected ROIs designated for invasive carcinoma based on the official pathology appraisal by 10x showed up-regulation of *ERBB2* (log FC = 0.97, -log10 P-adj = 54.8), *TACSTD2* (log FC = 1.09, -log10 P-adj = 50.6), and *EPCAM* (log FC = 1.06, -log10 P-adj = 42.9), which are highly expressed marker genes for breast cancers. This observation was supported by GO analysis, which revealed enrichment of the term ‘bicellular tight junction’, indicating the involvement of cancer epithelial cells in the ROIs. Additionally, cell type prediction analysis identified cancer epithelial cells as the most abundant cell type (37%) within the ROIs, further confirming the molecular profile of selected regions. Next, the ST of fresh frozen mouse colon tissue was analyzed with IAMSAM **(Figure 4b; Supplementary Figure 2)**. When we compared the segmentation results obtained from IAMSAM and the leiden clustering results, the segmentation results generated by IAMSAM showed remarkable agreement with the clustering of spots based on gene expression. In a specific example marked with a white arrow **(Figure 4b)**, the segmented region predominantly comprised stromal cells, aligning consistently with the corresponding GO terms associated with muscle and actin activities. When we applied the same analysis to the mouse brain coronal section, one of the most representative histological structures, the choroid plexus (CP), can be revealed in everything-mode **(Figure 4c; Supplementary Figure 3)**. The presence of cycling microglia as the predominant cell type along with CP cells supports the reasonable observation that border-associated macrophages, which is also a distinct cell type found in CP, were present.

We demonstrated the enhanced capabilities of the prompt-mode in IAMSAM through a case study involving ST-based analysis of tumor tissue from a 4T1-tumor bearing BALB/c mouse **(Figure 4d; Supplementary Figure 4)**. By utilizing the prompt-mode, we successfully defined an ROI in the stromal region located at the periphery of the tumor section (18), which was not achievable using the everything-mode. In this ROI, we identified chemokines secreted by M2 macrophages (e.g., *Cxcl12*) and stromal genes (e.g., *Fabp4*) among the top 10 expressed genes. Also, the high predicted abundance of Cd206 macrophages and fibroblasts within ROI supported the observations of the previous study (18). This case shows that the prompt-mode can offer enhanced flexibility in identifying ROI that the everything-mode may not capture.

To uncover microscopic histology features, the prompt-mode in IAMSAM can be particularly powerful, especially when used with magnification. In the Human_Prostate_Cancer sample, we demonstrated the capability of IAMSAM to identify and select microvessels as ROIs when utilizing the prompt-mode and the zoom-in interface **(Figure 4e; Supplementary Figure 5)**. Zooming-in on specific tissue areas helped identify microvessels, which may not be readily apparent on a larger scale. Furthermore, our analysis revealed that pan-endothelial cell markers, such as *CAV1* (log FC = 3.21, -log10 P-adj = 2.93), *CAV2* (log FC = 2.14, -log10 P-adj = 1.56), and *CAVIN1* (log FC = 1.50, - log 10 P-adj = 1.94) were up-regulated within the ROIs. In line with these findings, a GO term related to ‘focal adhesion’, specific to endothelial cells, was enriched, indicating the involvement of endothelial cells in these ROIs (21). Moreover, the cell type prediction analysis revealed that endothelial cells accounted for 27.7% of the total cell type population within the ROIs.

Lastly, we explored the application of IAMSAM with an optical image different from H&E staining **(Figure 4f; Supplementary Figure 6)**. We utilized a combined image of three distinct color channels corresponding to DAPI (4’,6-diamidino-2-phenylindole), Anti-GFAP, and Anti-NeuN staining. In this case, the successful identification of the Dentate Gyrus (DG) structure demonstrated the versatility and feasibility of IAMSAM in handling different imaging modalities. This finding further solidifies SAM as a general image segmentation algorithm that can be applied across various experimental setups. This feature highlights the broad applicability of IAMSAM and its potential to provide valuable information from diverse imaging modalities that spatially corresponds to ST data (18).

## Discussion

Integrating the Segment-Anything model into ST libraries has shown great promise as a new workflow to enhance the interpretation of image-based characteristics in ST data analysis, leading a more comprehensive molecular interpretation. This integration allows for accurate segmentation of histologically distinct structures and produces segments that align closely with gene expression clusters. By bridging the gap between histological features and molecular information, IAMSAM enables a comprehensive analysis of ST data. One of the strengths of IAMSAM lies in its user-friendly nature. This application has the potential to facilitate ST research by providing real-time and interactive tools for acquiring gene signatures from ST libraries. By lowering the barriers to entry in the field of ST, IAMSAM enables researchers to explore and analyze spatially resolved transcriptomic data more effectively. While IAMSAM offers powerful analysis capabilities, users must consider several factors to utilize the tool effectively.

First, it is crucial to select the proper mode when using IAMSAM. Our observations indicate that specific tissues may be better suited for the prompt-mode in IAMSAM, as this mode allows for capturing subtle and local differences that the everything-mode may overlook. As demonstrated by several use cases, we have significantly improved the segmentation performance by introducing an interactive function to IAMSAM. Second, selecting the appropriate reference model for cell type prediction is essential. IAMSAM utilizes the CellTypist API and offers multiple reference models. To select the model that best fits their sample, researchers should carefully consider variables like tissue type, species, and other pertinent characteristics. This decision guarantees reliable and precise cell-type predictions.

The spatial context of single-cell omics studies can be better understood by utilizing the spatial information offered by ST data, thereby providing futher insights into biology. Integrating this spatial information with diverse tissue images holds particular value, as it enhances the interpretability of the data (6). In this regard, exploring the implications of visually discernible histological characteristics, such as dense cancerous areas or stroma-rich regions on images, becomes crucial. The critical role of IAMSAM is to elucidate the molecular characteristics associated with these distinctive image regions and determine which cells exhibit such image-specific characteristics through the integration of ST data and image data analysis. In essence, this approach allows for a comprehensive understanding of the molecular attributes underlying visually distinctive patterns in the images and the specific cellular contributions to these patterns. Moreover, ST analysis effectively reveals heterogeneity based on tissue image characteristics, extending beyond a mere assessment of cellular composition. IAMSAM presents a workflow that explains visually identifiable image features with molecular information, offering a new direction for ST analysis.

## Conclusions

IAMSAM is a user-friendly web-based tool designed to analyze ST data. The tool utilizes the Segment-anything algorithm to segment H&E images of Visium data and performs statistical analysis to identify DEGs and their corresponding GO terms for each segmented region. With its simple and accessible interface, IAMSAM makes it easy for researchers to analyze and interpret their ST data. IAMSAM will be a valuable resource for researchers in the field of ST.

## Methods

### Dataset and preprocessing

We used six publicly accessible ST datasets, as shown in **Figure 4**, to illustrate the utility of IAMSAM. These datasets were chosen to represent a variety of tissues and experimental setups, allowing for a thorough assessment of IAMSAM’s capabilities. To exploit ST data for analysis in IAMSAM, the following five essential files, which are generated by the 10X Visium platform by default, should be organized as a folder within the ‘data’ folder: *‘filtered_feature_bc_matrix*.*h5’, ‘spatial/tissue_positions_list*.*csv’, ‘spatial/scalefactors_json*.*json’, ‘spatial/tissue_lowres_image*.*png’* and *‘spatial/tissue_hires_image*.*png’*. We employed the Scanpy package (22) to perform initial data manipulation steps for preprocessing. Specifically, spots containing fewer than 200 genes and genes with fewer than 200 counts were excluded from the feature matrix. Subsequently, a default log-normalization process was applied to each spot, facilitating the normalization and scaling of gene expression values across the dataset. Also, the pixel coordinates in the ST images were adjusted by multiplying them with the scale factor to align the image coordinates with the corresponding spot positions in the dataset. Additionally, the image was cropped to a smaller size that focused on the tissue area, excluding fiducial spots, while minimizing the padding around the image. This cropped image was used as input for the SAM.

### Segment-anything model

IAMSAM uses the SAM to segment tissue images derived from ST data. SAM enables users to define ROIs effortlessly by detecting morphologically distinct regions within the images. SAM consists of three components: an image encoder, a prompt encoder, and a mask decoder. The image encoder uses a pre-trained Vision Transformer (ViT) adapted to handle high-resolution inputs (23,24). This encoder runs once per image and can be applied before prompting the model. It encodes the input image into a high-dimensional vector that is then used as input for the mask decoder. The prompt encoder considers two types of prompts: sparse (points, boxes, text) and dense (masks). Sparse prompts include points and boxes, which are represented by positional encodings that are summed with learned embeddings for each prompt type (25).

In contrast, text prompts are handled differently compared to other sparse prompts. Instead of using positional encodings, text prompts are embedded using the CLIP framework (26). Dense prompts, which include masks, are embedded using convolutions and summed elementwise with the image embedding. The mask decoder maps the image embedding, prompt embeddings, and an output token to a mask. This component employs a modified Transformer decoder block and a dynamic mask prediction head (27). The decoder block uses prompt self-attention and cross-attention in two directions (prompt-to-image embedding and vice-versa) to update all embeddings. After running two blocks, the image embedding is upsampled, and an MLP maps the output token to a dynamic linear classifier, which computes the mask foreground probability at each image location. To address ambiguity in the prompt, the model is modified to predict multiple output masks for a single prompt, with each mask having a confidence score (estimated IoU) assigned to it.

### Two-modes of IAMSAM : everything-mode and prompt-mode

As SAM can perform segmentation incorporating manual prompts and automatic prompting, IAMSAM offers two main modes of operation: everything-mode and prompt-mode. In the everything-mode, the model performs image segmentation without any manual input from the user. It takes the input image and automatically generates segmentation masks for all different objects in the image. IAMSAM project segmentation masks the tissue image with a distinct colormap, enabling users to select ROI conveniently. When the user clicks on a mask, IAMSAM captures the coordinates of the click event and adds the corresponding mask to the list of selected regions.

On the other hand, the prompt-mode allows the user to provide additional information about the model. Users can draw multiple rectangles on the main visualization panel with *‘modeBarButtons*.*drawrect’*, the drawing tool in Plotly, by default. IAMSAM tracks the coordinates of rectangles and uses those as box prompts given to the SAM model. The model then generates the segmentation masks based on those input prompts, treating these masks as ROIs for further analysis.

### Mask filtering in everything mode

A mask filtering step was applied to remove unnecessary masks that do not contain any spots before visualizing the masks on the main visualization panel. Since the SAM model generates segmentation masks for the entire tissue image in the everything-mode, some masks may not have ST spots. To address this, a proportion-based filtering approach was implemented to retain only the masks that contain ST spots. The proportion of co-location between each mask and the spot coordinates was calculated by examining the overlap of mask pixels with the spot coordinates. If the calculated proportion was below 0.01, indicating a mask does not contain any spots, the corresponding mask was removed from the segmentation list. This filtering process ensured that only masks containing ST spots were retained for further analysis.

### Downstream analysis

We utilized various tools and packages to extract meaningful insights from the ST data in the downstream analysis step. To identify DEGs, we employed the *‘sc*.*tl*.*rank_genes_groups’* function in Scanpy, employing the wilcoxon method. This analysis allows for the calculation of statistical significance and enables the identification of genes that exhibit significant differences in expression between conditions or cell types. Users have the flexibility to modify the cutoff values for adjusted p-value and log fold changes, enabling manual DEG definition. We specifically focus on displaying the top 10 genes with the biggest fold changes for the box plots representing DEGs. Enrichment analysis uses the ‘enrichr’ function from the GSEApy package (28). In addition to DEG and enrichment analyses, IAMSAM incorporates cell type predictions using the CellTypist API (13). The probability matrix from the CellTypist result is normalized by dividing each row by the sum of its values, representing the relative likelihood of each cell belonging to a specific cell type. The probabilities for each cell type are summed up, and IAMSAM presents the top six cell types with the highest proportions in a pie chart. All visualizations in IAMSAM are created using the Plotly package.

### User interface and web application

IAMSAM is built as a Dash application, leveraging its powerful framework for creating interactive web-based data visualization and analysis tools (12). The Dash framework, built on top of Flask, Plotly, and React, provides a highly customizable and responsive user interface. The user interface of IAMSAM is designed to provide a seamless and intuitive experience for researchers analyzing ST data. The main components of the user interface include a visualization panel with a dropdown menu to select samples to analyze, parameter panels that affect the SAM model and analysis result, and downstream analysis panels. IAMSAM offers two main modes, everything-mode, and prompt-mode, accessible through a tab menu. This allows users to easily navigate between the modes. Being a web application, IAMSAM takes advantage of various interactive features to enhance user interaction and data exploration. Users can dynamically adjust parameters, such as fold change and p-value cutoff, to customize the analysis results. The visualizations, such as volcano plots and box plots, are interactive and allow users to zoom in, zoom out, and capture specific ROI for further examination.

## Acknowledgements

We would like to express our sincere gratitude to Meta AI for their invaluable contribution in making the source code of Segment-anything available to the public. Their unwavering commitment to the open-community ethos has not only greatly facilitated our research but has also empowered researchers and developers worldwide to explore and innovate in the field of image segmentation. We would like to express our gratitude to all the researchers at Portrai, Inc. for their support and contributions.

## Availability of data and materials

The datasets used in this study are publicly available and can be accessed through the following sources. The human breast cancer and prostate cancer data obtained by the Visium FFPE protocol can be downloaded from the 10X Genomics public repository (https://www.10xgenomics.com/resources/datasets/). Mouse brain coronal sections stained with DAPI and Anti-NeuN can also be obtained from the 10X Genomics. The datasets for mouse colon, mouse coronal brain section, and mouse syngeneic 4T1 tumor data are available from the Gene Expression Omnibus (GEO) under the accession number GSM5213483, GSM5519060, and GSE196506, respectively. The code for the IAMSAM is publicly available at the GitHub repository (https://github.com/portrai-io/IAMSAM).

## Notes

### Competing Interest Statement

The authors have declared no competing interest.

