## Supplementary Figures for "IAMSAM : Image-based Analysis of Molecular signatures using the Segment-Anything Model"

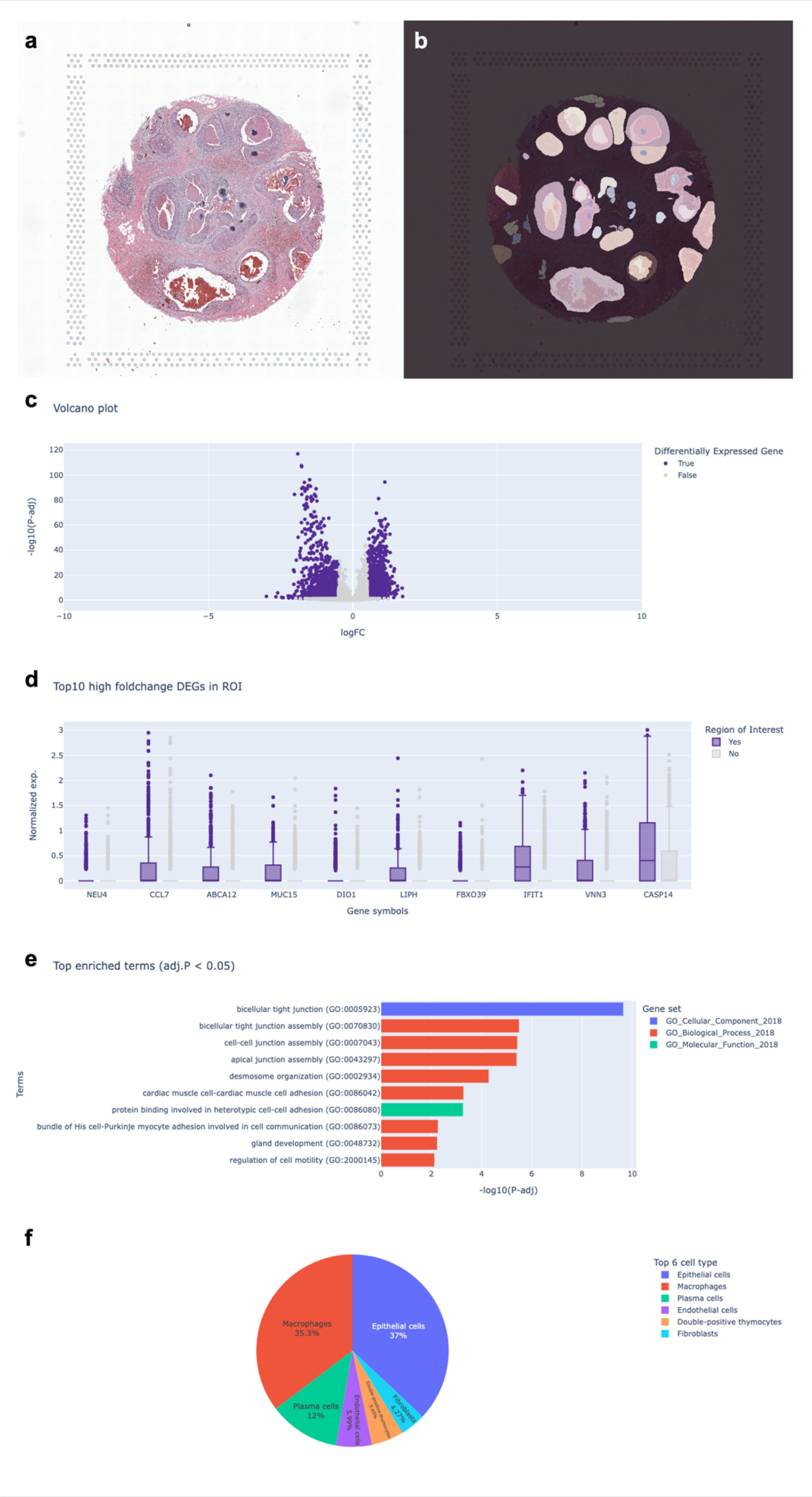
a
b
c
Volcano plot
Differentially Expressed Gene
True
False
-log10(P-adj)
logFC
d
Top10 high foldchange DEGs in ROI
Region of Interest
Yes
No
Normalized exp.
NEU4
CCL7
ABCA12
MUC15
DIO1
LIPH
FBXO39
IFIT1
VNN3
CASP14
Gene symbols
e
Top enriched terms (adj.P < 0.05)
Gene set
GO_Cellular_Component_2018
GO_Biological_Process_2018
GO_Molecular_Function_2018
Terms
bicellular tight junction (GO:0005923)
bicellular tight junction assembly (GO:0070830)
cell-cell junction assembly (GO:0007043)
apical junction assembly (GO:0043297)
desmosome organization (GO:0002934)
cardiac muscle cell-cardiac muscle cell adhesion (GO:0086042)
protein binding involved in heterotypic cell-cell adhesion (GO:0086080)
bundle of His cell-Purkinje myocyte adhesion involved in cell communication (GO:0086073)
gland development (GO:0048732)
regulation of cell motility (GO:2000145)
-log10(P-adj)
f
Top 6 cell type
Epithelial cells
Macrophages
Plasma cells
Endothelial cells
Double-positive thymocytes
Fibroblasts
Macrophages 35.3%
Epithelial cells 37%
Plasma cells 12%
Endothelial cells 5.99%
Fibroblasts 4.27%

**Supplementary Figure 1 |** Analysis of invasive carcinoma regions in the FFPE slice of human breast cancer from 10x Genomics (a) The H&E-stained image from the ST data. (b) ROIs are selected using everything-mode with an IOU score threshold of 0.9. The regions identified for invasive carcinoma were determined based on the formal pathology evaluation. The following analyses include : (c) a volcano plot with a log FC threshold of 0.5 and an adjusted p-value threshold 0.05, (d) top 10 high genes in the ROIs (adjusted p-value < 0.05; ordered by log FC) compared with other regions in the form of box plots, (e) top enriched GO terms for all the up-regulated DEGs in the ROIs and (f) a pie chart showing cell type proportions using Immune_All_High.pkl reference model in CellTypist. Here, breast cancer-specific genes, including ERBB2, TACSTD2, and EPCAM, were up-regulated in the ROI as expected. Additionally, the analysis using CellTypist revealed that cancer epithelial cells were the most abundant cell type in the selected regions.

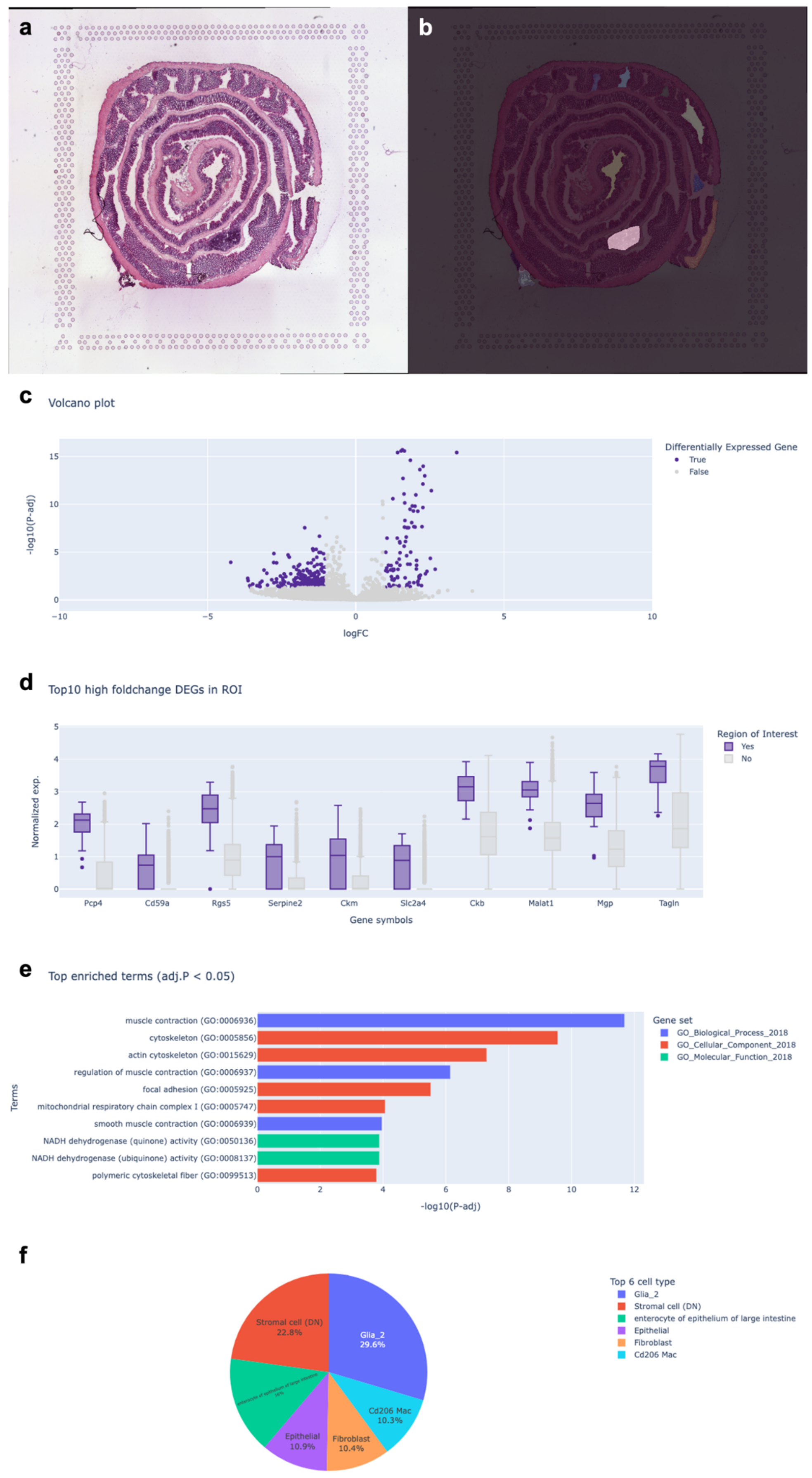
a
b
c
Volcano plot
Differentially Expressed Gene
True
False
-log10(P-adj)
logFC
d
Top10 high foldchange DEGs in ROI
Region of Interest
Yes
No
Normalized exp.
Pcp4
Cd59a
Rgs5
Serpine2
Ckm
Slc2a4
Ckb
Malat1
Mgp
Tagln
Gene symbols
e
Top enriched terms (adj.P < 0.05)
Gene set
GO_Biological_Process_2018
GO_Cellular_Component_2018
GO_Molecular_Function_2018
Terms
muscle contraction (GO:0006936)
cytoskeleton (GO:0005856)
actin cytoskeleton (GO:0015629)
regulation of muscle contraction (GO:0006937)
focal adhesion (GO:0005925)
mitochondrial respiratory chain complex I (GO:0005747)
smooth muscle contraction (GO:0006939)
NADH dehydrogenase (quinone) activity (GO:0050136)
NADH dehydrogenase (ubiquinone) activity (GO:0008137)
polymeric cytoskeletal fiber (GO:0099513)
-log10(P-adj)
f
Top 6 cell type
Glia_2
Stromal cell (DN)
enterocyte of epithelium of large intestine
Epithelial
Fibroblast
Cd206 Mac
Stromal cell (DN) 22.8%
Glia_2 29.6%
Cd206 Mac 10.3%
Fibroblast 10.4%
Epithelial 10.9%

**Supplementary Figure 2 |** Analysis of fresh frozen colon tissue from wild C57BL/6J mouse. (a) The H&E-stained image from the ST data (b) ROIs selected using the everything-mode with an IOU score threshold of 0.95. The chosen region aligned well with a spatial cluster from gene expression-based spot clustering in scanpy. The following analyses include: (c) a volcano plot with logFC threshold of 1 and adjusted p-value threshold of 0.05, (d) top 10 high genes in the ROIs (adjusted p-value < 0.05; ordered by logFC) compared with other regions in the form of box plots , (e) top enriched GO terms for all the up-regulated DEGs in the ROIs, and (f) a pie chart showing cell type proportions using Adult_Mouse_Gut.pkl reference model in CellTypist. The presence of stromal cells is consistent with the GO terms indicating muscle and actin activities.

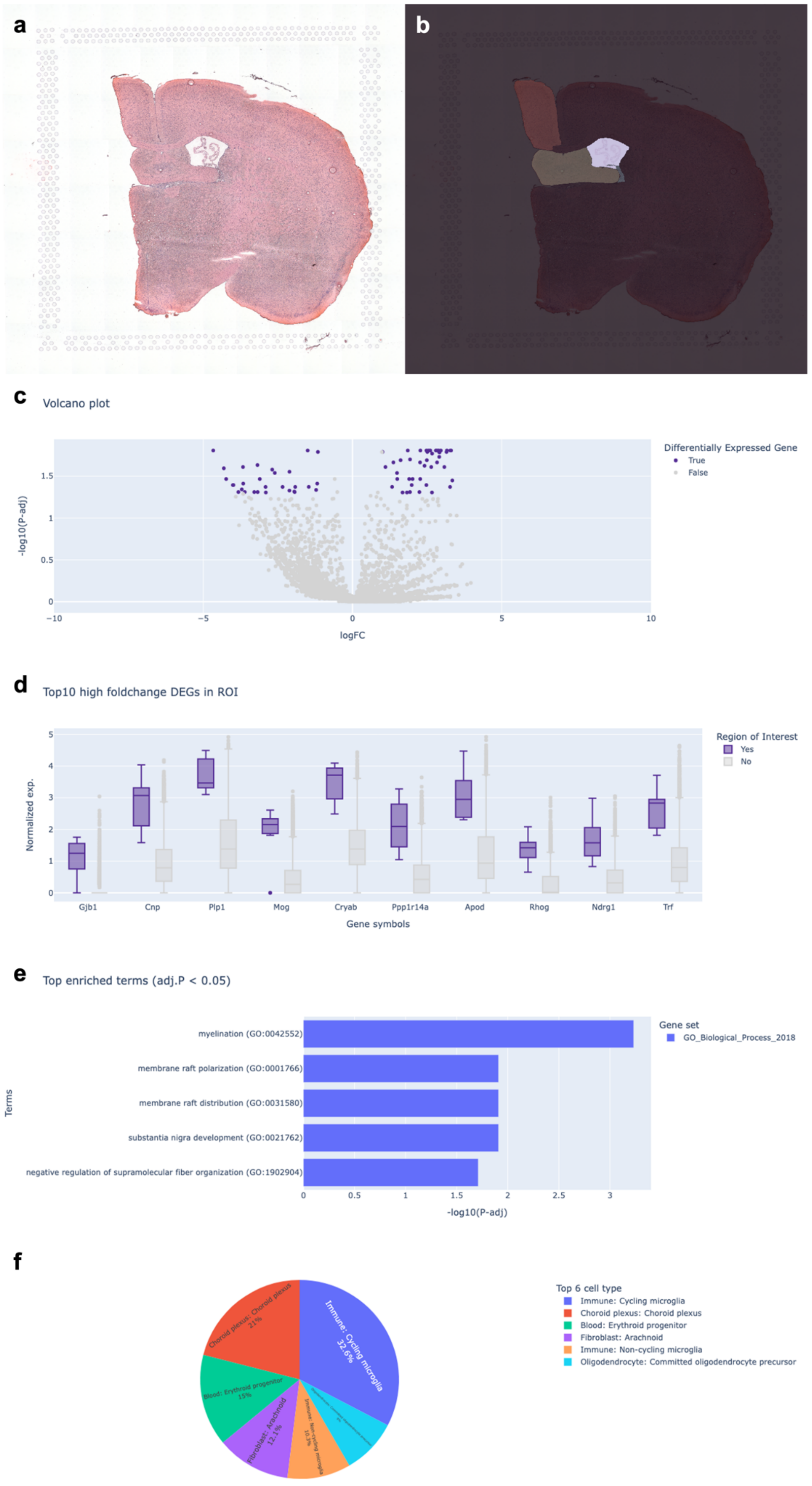
a
b
c
Volcano plot
-log10(P-adj)
1.5
1
0.5
0
−10
−5
0
5
10
logFC
Differentially Expressed Gene
True
False
d
Top10 high foldchange DEGs in ROI
Normalized exp.
5
4
3
2
1
0
Gjb1
Cnp
Plp1
Mog
Cryab
Ppp1r14a
Apod
Rhog
Ndrg1
Trf
Gene symbols
Region of Interest
Yes
No
e
Top enriched terms (adj.P < 0.05)
Terms
myelination (GO:0042552)
membrane raft polarization (GO:0001766)
membrane raft distribution (GO:0031580)
substantia nigra development (GO:0021762)
negative regulation of supramolecular fiber organization (GO:1902904)
0
0.5
1
1.5
2
2.5
3
-log10(P-adj)
Gene set
GO_Biological_Process_2018
f
Immune: Cycling microglia
32.6%
Choroid plexus: Choroid plexus
21%
Blood: Erythroid progenitor
15%
Fibroblast: Arachnoid
12.1%
Immune: Non-cycling microglia
10.3%
Top 6 cell type
Immune: Cycling microglia
Choroid plexus: Choroid plexus
Blood: Erythroid progenitor
Fibroblast: Arachnoid
Immune: Non-cycling microglia
Oligodendrocyte: Committed oligodendrocyte precursor

**Supplementary Figure 3 |** Analysis of a coronal section of a fresh frozen brain from a male C57BL/6J mouse. (a) The H&E stained image from ST data (b) ROIs selected using everything-mode with an IOU score threshold of 0.95. The chosen region well represented the choroid plexus structure of a brain. The following analyses include: (c) a volcano plot with logFC threshold of 1 and adjusted p-value threshold of 0.05, (d) top 10 high genes in the ROIs (adjusted p-value < 0.05; ordered by logFC) compared with other regions in the form of box plots, (e) top enriched GO terms for all the up-regulated DEGs in the ROIs, and (f) a pie chart showing cell type proportions using Developing_Mouse_Brain.pkl reference model in CellTypist (f). Here, the choroid plexus (CP) appeared as expected.

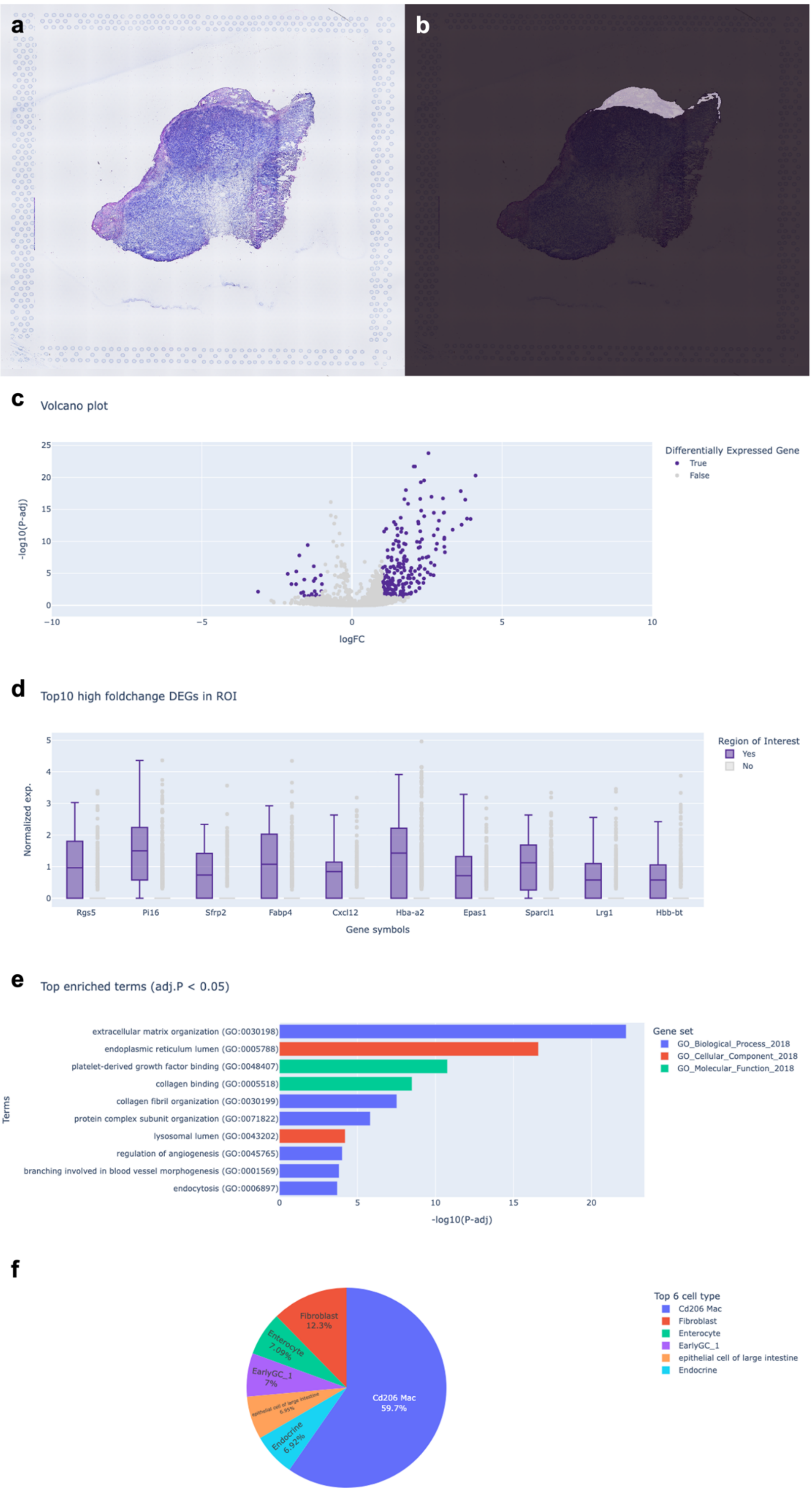
a
b
c
Volcano plot
Differentially Expressed Gene
True
False
-log10(P-adj)
logFC
d
Top10 high foldchange DEGs in ROI
Region of Interest
Yes
No
Normalized exp.
Rgs5
Pi16
Sfrp2
Fabp4
Cxcl12
Hba-a2
Epas1
Sparcl1
Lrg1
Hbb-bt
Gene symbols
e
Top enriched terms (adj.P < 0.05)
extracellular matrix organization (GO:0030198)
endoplasmic reticulum lumen (GO:0005788)
platelet-derived growth factor binding (GO:0048407)
collagen binding (GO:0005518)
collagen fibril organization (GO:0030199)
protein complex subunit organization (GO:0071822)
lysosomal lumen (GO:0043202)
regulation of angiogenesis (GO:0045765)
branching involved in blood vessel morphogenesis (GO:0001569)
endocytosis (GO:0006897)
Terms
-log10(P-adj)
Gene set
GO_Biological_Process_2018
GO_Cellular_Component_2018
GO_Molecular_Function_2018
f
Top 6 cell type
Cd206 Mac
Fibroblast
Enterocyte
EarlyGC_1
epithelial cell of large intestine
Endocrine
Cd206 Mac 59.7%
Fibroblast 12.3%
Enterocyte 7.09%
EarlyGC_1 7%
Endocrine 6.92%

**Supplementary Figure 4 |** Analysis of fresh frozen section of a 4T1 syngeneic tumor from a female BALB/c mouse. (a) The H&E stained image from ST data. (b) ROIs selected for a tumor stromal region. The following analyses include: (c) a volcano plot with logFC threshold of 1 and adjusted p-value threshold of 0.05, (d) top 10 high genes in the ROIs (adjusted p-value < 0.05; ordered by logFC) compared with other regions in the form of box plots, (e) top enriched GO terms for all the up-regulated DEGs in the ROIs, and (f) a pie chart showing cell type proportions using Adult_Mouse_Gut.pkl reference model in CellTypist (f).It displays the proportions of Cd206 macrophages (M2 macrophages) and fibroblasts, which are predominant in the stromal region analyzed.

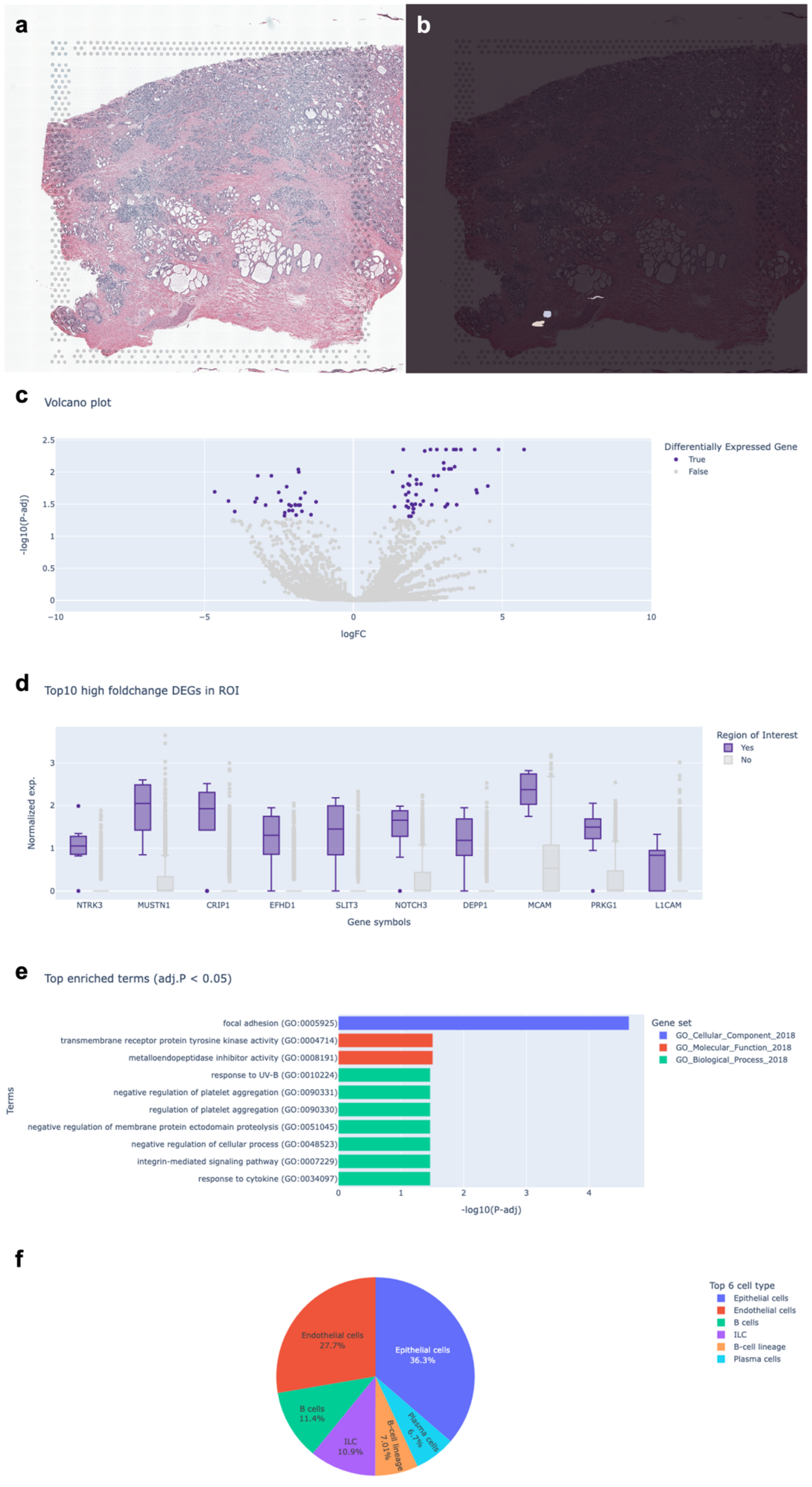
a
b
c
Volcano plot
-log10(P-adj)
logFC
Differentially Expressed Gene
True
False
d
Top10 high foldchange DEGs in ROI
Normalized exp.
Gene symbols
NTRK3
MUSTN1
CRIP1
EFHD1
SLIT3
NOTCH3
DEPP1
MCAM
PRKG1
L1CAM
Region of Interest
Yes
No
e
Top enriched terms (adj.P < 0.05)
Terms
focal adhesion (GO:0005925)
transmembrane receptor protein tyrosine kinase activity (GO:0004714)
metalloendopeptidase inhibitor activity (GO:0008191)
response to UV-B (GO:0010224)
negative regulation of platelet aggregation (GO:0090331)
regulation of platelet aggregation (GO:0090330)
negative regulation of membrane protein ectodomain proteolysis (GO:0051045)
negative regulation of cellular process (GO:0048523)
integrin-mediated signaling pathway (GO:0007229)
response to cytokine (GO:0034097)
-log10(P-adj)
Gene set
GO_Cellular_Component_2018
GO_Molecular_Function_2018
GO_Biological_Process_2018
f
Endothelial cells 27.7%
Epithelial cells 36.3%
B cells 11.4%
ILC 10.9%
B-cell lineage 7.01%
Plasma cells 6.7%
Top 6 cell type
Epithelial cells
Endothelial cells
B cells
ILC
B-cell lineage
Plasma cells

**Supplementary Figure 5 |** Analysis of vessels in an FFPE slice of human prostate cancer from 10X Genomics. (a) The H&E stained image from ST data. (b) The selection of vessels was performed using the prompt-mode in conjunction with a zoom-in interface, with validation from a pathologist to ensure accurate identification. The following analyses include: (c) a volcano plot with logFC threshold of 1 and adjusted p-value threshold of 0.05, (d) top 10 high genes in the ROIs (adjusted p-value < 0.05; ordered by log FC) compared with other regions in the form of box plots, (e) top enriched GO terms for all the up-regulated DEGs in the ROIs, and (f) a pie chart showing cell type proportions using Immune_All_High.pkl reference model in CellTypist. Here, endothelial cells occupying 27.7% of the total cell type population were significantly abundant compared to the ROIs.

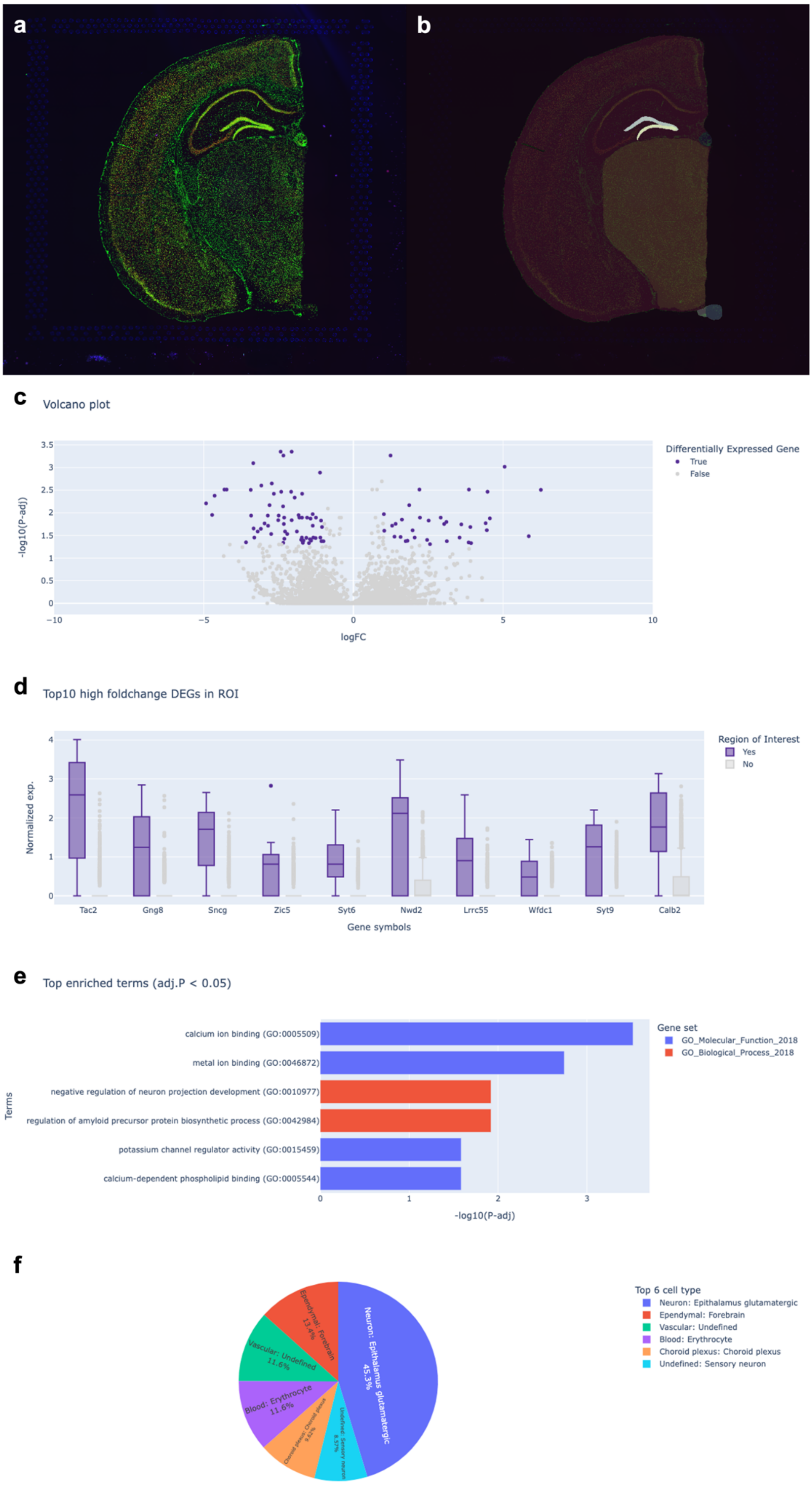
a
b
c
Volcano plot
Differentially Expressed Gene
True
False
-log10(P-adj)
logFC
d
Top10 high foldchange DEGs in ROI
Region of Interest
Yes
No
Normalized exp.
Tac2
Gng8
Sncg
Zic5
Syt6
Nwd2
Lrrc55
Wfdc1
Syt9
Calb2
Gene symbols
e
Top enriched terms (adj.P < 0.05)
Gene set
GO_Molecular_Function_2018
GO_Biological_Process_2018
calcium ion binding (GO:0005509)
metal ion binding (GO:0046872)
negative regulation of neuron projection development (GO:0010977)
regulation of amyloid precursor protein biosynthetic process (GO:0042984)
potassium channel regulator activity (GO:0015459)
calcium-dependent phospholipid binding (GO:0005544)
Terms
-log10(P-adj)
f
Top 6 cell type
Neuron: Epithalamus glutamatergic
Ependymal: Forebrain
Vascular: Undefined
Blood: Erythrocyte
Choroid plexus: Choroid plexus
Undefined: Sensory neuron
Neuron: Epithalamus glutamatergic 45.3%
Ependymal: Forebrain 13.4%
Vascular: Undefined 11.6%
Blood: Erythrocyte 11.6%

**Supplementary Figure 6 |** Analysis of a coronal section of a mouse brain with fluorescence image (a) The three-channel fluorescence image from ST data (b) The selection of the Dentatus Gyrus (DG) by using everything-mode with IOU score threshold of 0.85. The following analyses including a volcano plot with logFC threshold of 1 and adjusted p-value threshold of 0.05 (c), top 10 high genes in the ROIs (adjusted p-value < 0.05; ordered by logFC) compared with other regions in the form of box plots (d), top enriched GO terms for all the up-regulated DEGs in the ROIs (e), and a pie chart showing cell type proportions using Developing_Mouse_Brain.pkl reference model in CellTypist (f).
